# Enrichment of the HIV reservoir in CD32+ CD4 T cells occurs early in blood and tissue

**DOI:** 10.1101/169342

**Authors:** Genevieve E Martin, Matthew Pace, John P Thornhill, Chansavath Phetsouphanh, Jodi Meyerowitz, Morgane Gossez, Emily Hopkins, Helen Brown, Nicola Robinson, Natalia Olejniczak, Gita Ramjee, Pontiano Kaleebu, Kholoud Porter, Christian Willberg, Paul Klenerman, Nneka Nwokolo, Julie Fox, Sarah Fidler, John Frater on behalf of the CHERUB investigators

## Abstract

The Fc receptor CD32 has been proposed as a marker for CD4 T cells latently infected with HIV. We demonstrate that enrichment for HIV DNA in CD32+ CD4 T cells can be found early in infection in both tissue and blood. However, we find no evidence for a correlation between CD32 expression on CD4 T cells and either HIV DNA levels or time to rebound viraemia following treatment interruption. CD32+ CD4 T cells have a more differentiated memory phenotype, and high levels of expression of immune checkpoint receptors PD-1, Tim-3 and TIGIT as well as the activation marker, HLA DR. There was no difference in the phenotype or frequency of CD32 expressing cells prior to or after the initiation of antiretroviral therapy, or compared with healthy controls, suggesting that preferential infection or survival, rather than up-regulation, may be responsible for the observed enrichment of proviral HIV DNA in CD32+ CD4 T cells.

## Introduction

A cure for HIV infection is contingent on targeting and neutralising a reservoir of latently infected cells that persist despite effective antiretroviral therapy. These cells are predominantly resting memory CD4+ T cells and contain transcriptionally-silent integrated proviral HIV DNA. Activation of these cells results in HIV production and disease progression in the absence of ART. Latently infected cells are extremely rare (0.0001 - 0.1% of CD4 T cells), complicating their study. Accordingly, there has been a research effort to identify cell surface markers which identify latently infected cells to facilitate their characterisation and targeting. For example, immune checkpoint receptors (ICRs)^1^ such as Programmed Death Receptor-1 (PD-1), T cell immunoglobulin and mucin domain-containing molecule (Tim)-3 and T cell immunoreceptor with immunoglobulin and ITIM domains (TIGIT) are associated with an exhausted effector state^2-5^ and their expression correlates with HIV disease progression and viral rebound following treatment interruption^4, 6, 7^. ICRs have been reported to be highly expressed on cells comprising the HIV reservoir^8, 9^, but are not discriminatory. Equally, CD2 was reported to be enriched on latently infected cells, but is also widely expressed on other T cells^10^.

Recently, CD32a (FcγRIIa) - a low affinity IgG receptor expressed on myeloid cells and granulocytes, but not generally considered to be expressed on T cells^11^ – has been proposed as a specific surface marker for latent HIV infection^12^. It is unclear whether this enrichment holds true in tissue^13^ as well as peripheral blood CD4 T cells, or whether there is differential expression in resting memory cells which comprise a major part of the latent reservoir^9, 14, 15^. Here we investigate CD32 expression on CD3+ CD4+ cells in the blood and tissue of individuals treated during primary HIV infection (PHI) (a group of interest due to an association with post treatment virological remission and a more labile reservoir^16-18^), and explore associations with cell phenotype, other putative reservoir markers and clinical progression.

## Results

### Participants

We studied individuals who commenced antiretroviral therapy (ART) during primary HIV infection (PHI) from two clinical trials (SPARTAC^19^ and HEATHER^7^). Key demographic and clinical characteristics are shown in Table 1. The median time from estimated seroconversion to ART initiation was 74 days (interquartile range, IQR 36 – 114 days). Median baseline plasma viral load (pVL) was 4.94 log_10_ [copies/ml] (IQR 4.09 – 6.52) and CD4 T cell count was 545 cells/μl (IQR 453 - 696). Clinical, immunological and reservoir measures presented are at baseline (before the initiation of ART) and after 1 year of ART (median 48.1 weeks; IQR 47.9 to 52.9 weeks). Healthy controls (n=10; 100% male; median age 34.5 [IQR 30.5-42.5] years) were included for comparison.

**Table 1.** Demographic and clinical characteristics of participants. Demographic and clinical characteristics of participants included in main studies. Values given represent n (%) for categorical variables and median (interquartile range) for continuous variables.

|  | HEATHER (n=20) | SPARTAC (n=19) |
| --- | --- | --- |
| Sex |  |  |
| • Male | 20 (100%) | 5 (26%) |
| Age | 33 (28 – 40.8) | 27 (22 – 40) |
| Country |  |  |
| • United Kingdom | 20 (100%) | 4 (21%) |
| • South Africa/Uganda | 0 | 13 (68%) |
| • Other | 0 | 2 (11%) |
| Time between estimated date of seroconversion and ART initiation (days) | 41.5 (22.5 – 55.3) | 109 (76 – 124) |
| Time of sampling (weeks since ART initiation) | 52.5 (52.0 – 57.6) | 47.9 (47.6 – 48.0) |
| Baseline CD4 T cell count (cells/μL) | 514 (376 – 628) | 634 (535 – 764) |
| Baseline HIV RNA ($\log_{10}$ copies/mL) | 6.32 (4.55 – 6.76) | 4.32 (3.68 – 4.95) |

### *CD32* + CD4 T cells are enriched for HIV DNA

For participants on ART, the median percentage of CD4 T cells expressing CD32 was 1.5% (range 0.24–6.4%; representative gating shown in Fig. 1a; further examples in Supplementary Fig. 1). This did not differ from pre-therapy levels (median 1.4%; range 0.31-2.6%) or healthy controls (median 1.7%; range 0.49–4.9%; Fig. 1b; p=0.55). In treated individuals (n=6, Supplementary Table 1), sorted CD32+ CD4 T cells were highly enriched for HIV DNA compared with CD32-cells (median 69-fold enrichment; range 16-333 fold; Fig 1c), providing the first confirmation of CD32 as a marker of an enriched reservoir. The percentage of CD32+ CD4 T cells, however, did not correlate with reservoir size as measured by HIV DNA in CD4 T cells either on ART (Fig. 1d) or prior to ART initiation (Supplementary Fig. 2). The fold enrichment varied significantly between individuals, which may partially explain the observed absence of correlation.

**Figure 1.**
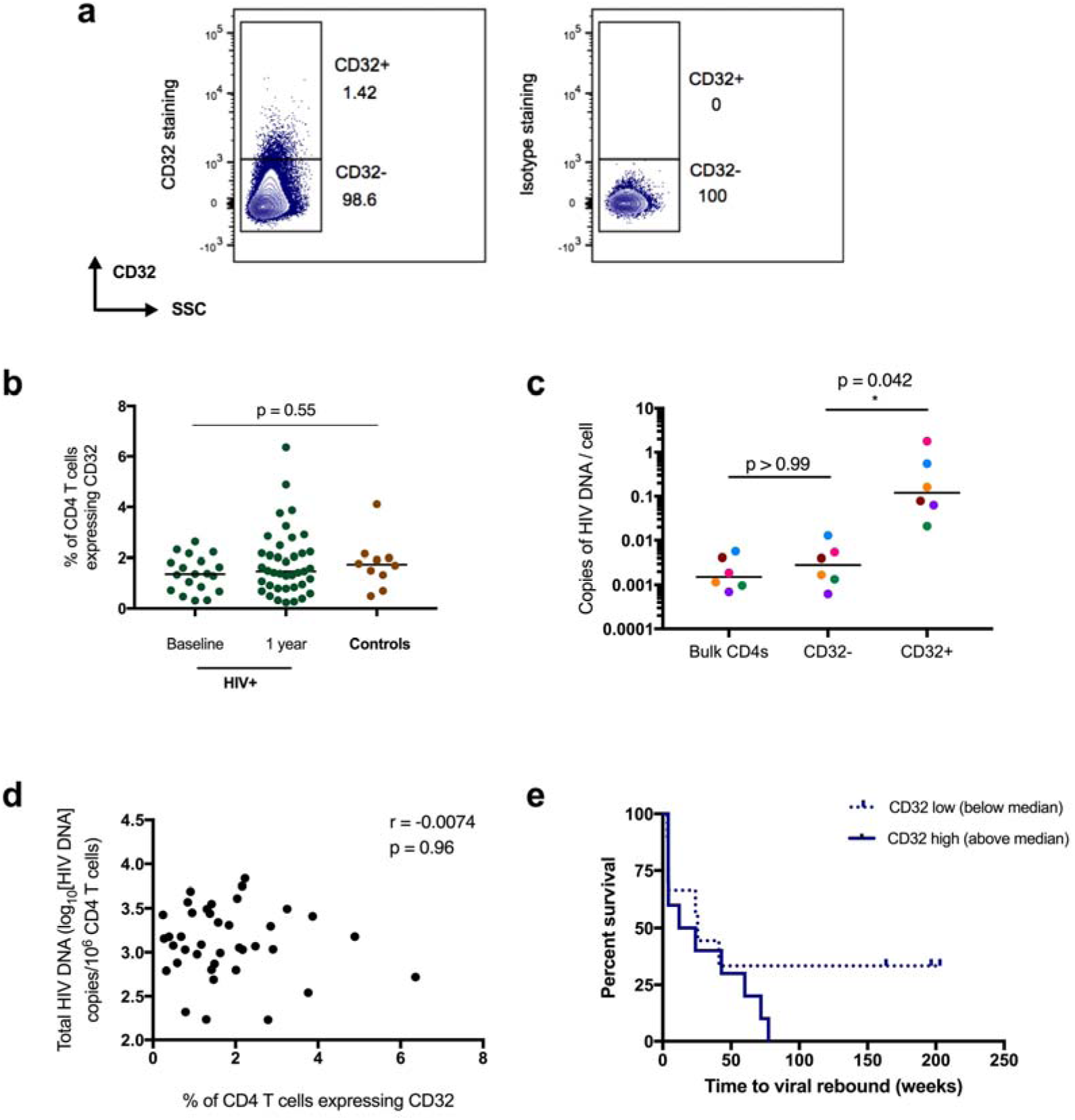
CD32 expression and HIV reservoir quantitation. Representative gating of CD32 on CD4 T cells from an HIV+ individual is shown (left panel) in (a), relative to isotype control (right panel). The percentage of CD32+ CD4 T cells is shown in (b) for HIV+ individuals with primary HIV infection pre-therapy (baseline; n=19) and 1 year following ART initiation (n=39), as well as for controls (n=10); groups were compared using a Kruskal-Wallis test. Panel (c) shows HIV DNA in bulk CD4s as well as sorted CD32+ and CD32-CD4 T cells from HIV+ individuals (n=6) at 1 year post-ART initiation. All three groups were compared using a Friedman test (p=0.0055) with the p-values shown corresponding to subsequent pairwise comparisons with Dunn’s multiple comparison test. For (b) and (c) bars indicate the median value. The relationship between HIV DNA and percentage of CD32+ CD4 T cells in treated HIV+ individuals (n=39), shown in (d), was assessed using Spearman’s rank correlation. Panel (e) shows survival curves (Kaplan-Meier) of time to viral rebound >400 copies/mL based on CD32 expression (n=19) stratified at the median. All bars shown indicate the median value; ∗ indicates p-value <0.05.

Whether percentage of CD32+ CD4 T cells predicted time to pVL rebound after ART treatment interruption (TI) was assessed in a subset of individuals after 48 weeks of suppressive ART (n=19). In Cox models, we found no evidence to support such an association for CD32 expression (HR 1.2 (95% CI 0.85 – 1.8), p=0.29). There was, however, weak evidence of an association between time to rebound and baseline pVL (p=0.053), but not with baseline CD4 T cell count or HIV DNA at TI, parameters which had been associated with time to rebound in a large sub-analysis of the SPARTAC^20^ (Supplementary Table 2). Three individuals remained virologically suppressed at the end of follow up after TI (median 197 weeks), all of whom had CD32+ CD4% values below the median (Fig. 1e).

### CD32 expression on CD4 T cells is associated with a differentiated memory phenotype

As CD4 T cells have not previously been considered to express CD32, little is known about which CD4 T cells express this marker. We therefore compared the memory phenotype of CD32+ and CD32-CD4 T cell subsets (as defined in Fig. 2a) during treated PHI. CD32+ CD4 T cells had lower proportions of naÏve and central memory (T_CM_) T cells than their CD32-counterparts. In contrast, transitional memory (T_TM_), effector memory (T_EM_) and T_EMRA_ cells comprised a greater portion of the CD32+ CD4 T cell pool than for CD32-CD4 T cells (Fig. 2b; all p<0.001), reflecting a more differentiated memory phenotype. The same pattern was also observed in healthy controls (Fig. 2c) and prior to ART initiation (Supplementary Fig. 3).

**Figure 2.**
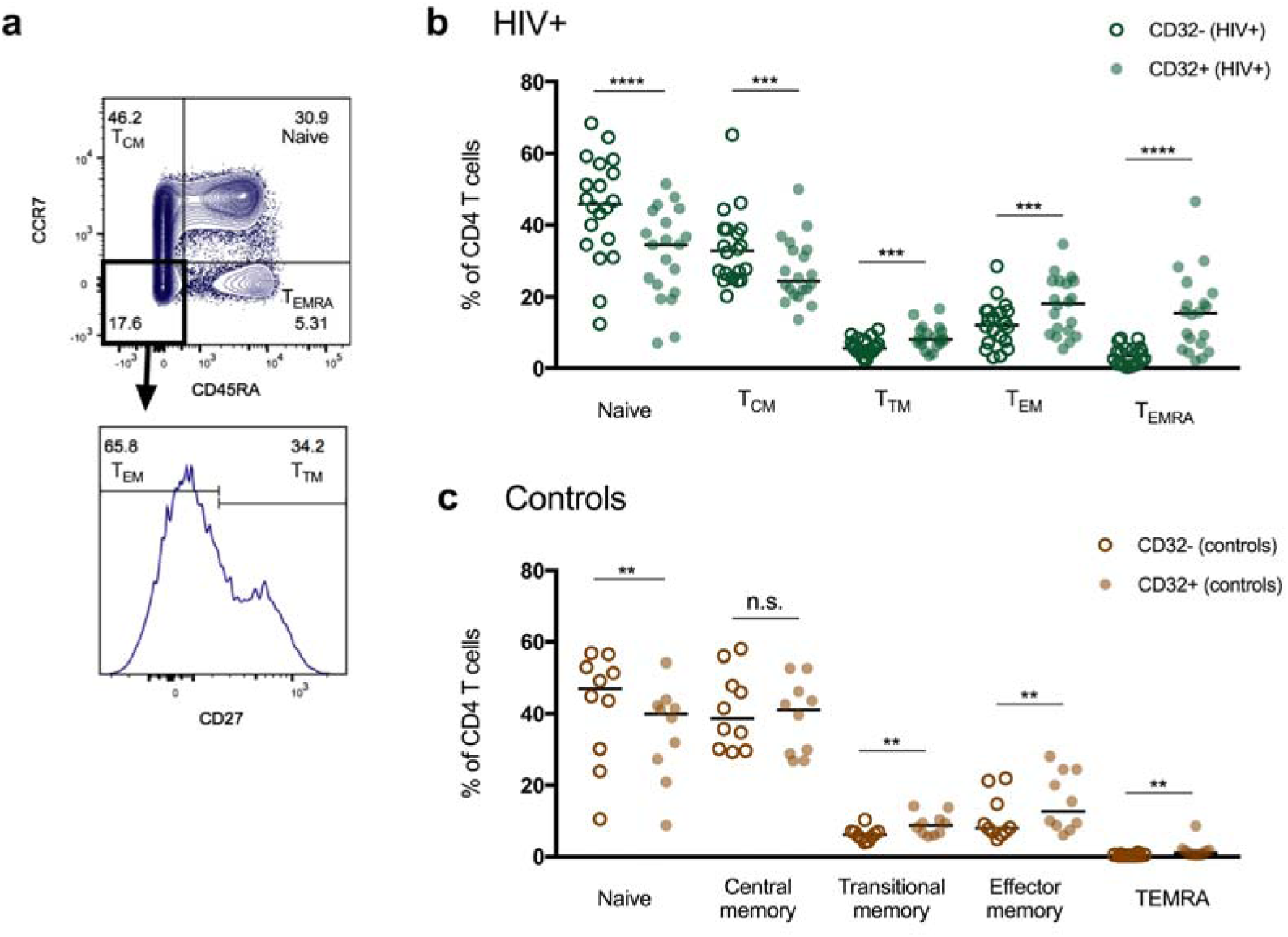
Memory differentiation of CD32+ and CD32-CD4 T cells. The percentage of CD32+ and CD32-CD4 T cells comprised of naïve, central memory (T_CM_), transitional memory (T_TM_), effector memory (T_EM_) and T_EMRA_ cells was quantified by flow cytometry based on the expression of CD45RA, CCR7 and CD27. Representative gating is shown in (a). Panel (b) shows the difference in memory distribution between CD32+ (closed circles) and CD32-(open circles) CD4 T cells in HIV+ individuals (n=20) at 1 year following the initiation of antiretroviral therapy. CD32+ and CD32-subsets were compared using the Wilcoxon matched-pairs signed rank test. Bars indicate the median value. The same analysis is shown for healthy controls (n=10) in (c). ∗∗∗∗ indicates p<0.0001, ∗∗∗ indicates p 0.0001-0.001, ∗∗ indicates p 0.001-0.01, n.s. (non-significant) indicates p≥0.05.

### CD32 expressing CD4 T cells have high levels of immune checkpoint receptors

The expression of PD-1, Tim-3 and TIGIT was elevated on CD32+ CD4 T cells compared with CD32-counterparts, in both treated HIV+ individuals and healthy controls (Figure 3a-d), although without significant correlations between ICR and CD32 expression (Supplementary Fig. 4). Elevated expression of all three ICRs on CD32+ CD4 T cells was also observed at baseline (Supplementary Fig. 5) and, for PD-1 and Tim-3, at higher levels than after a year of ART. These ICRs are known to be variably expressed on different CD4 memory subsets, with elevated expression on non-naïve T cells^8, 9, 21^. Elevated PD-1, Tim-3 and TIGIT expression was observed even when accounting for memory differentiation (Supplementary Fig 6), excluding this as a confounder for the observed elevated ICR expression.

**Figure 3.**
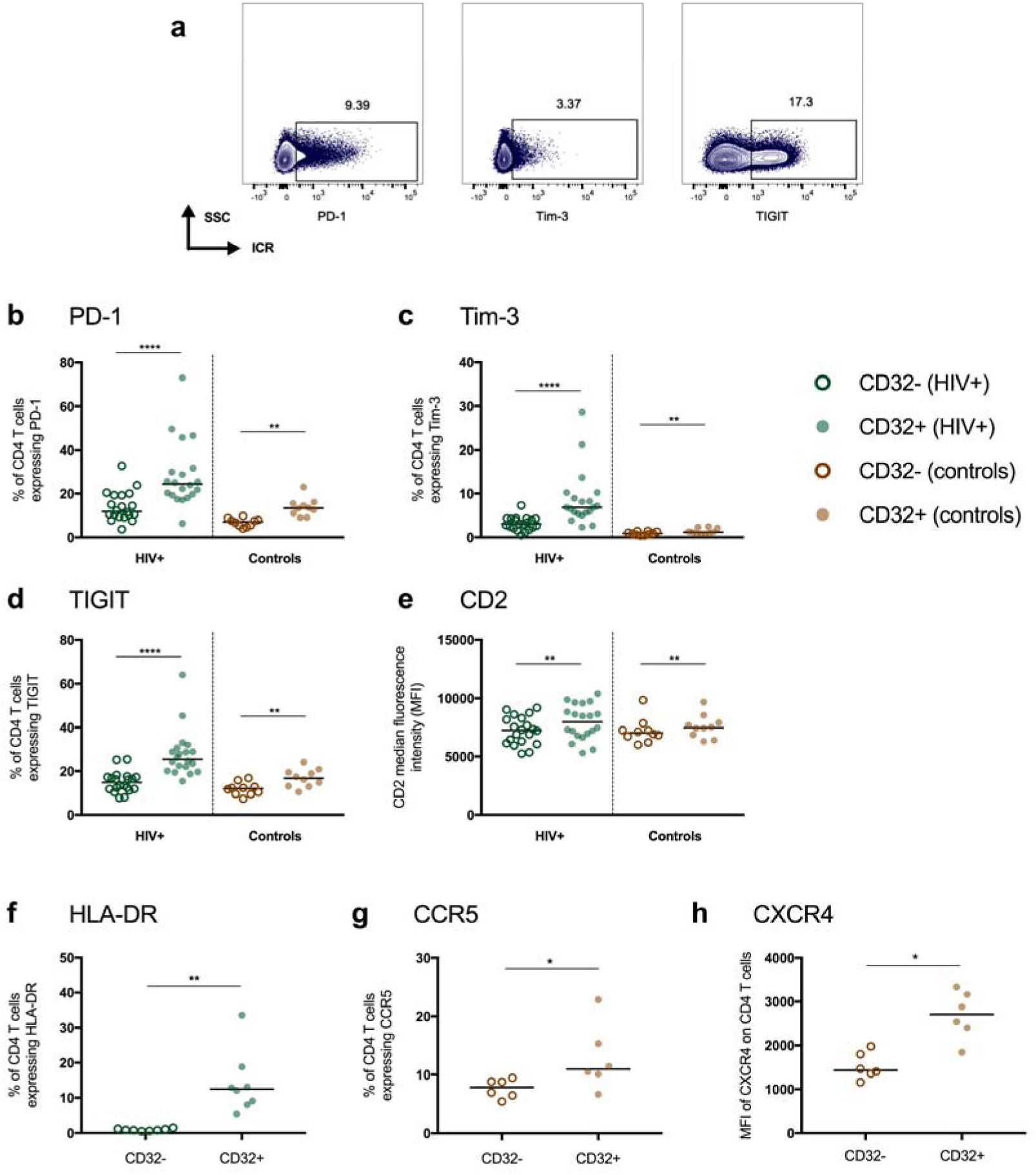
Immune checkpoint receptor, activation marker and CD2 expression on peripheral blood CD32+ and CD32-CD4 T cells. The expression of PD-1, Tim-3 and TIGIT was measured on CD4 T cell subsets by flow cytometry, with representative gating shown in (a). The expression of PD-1 (b), Tim-3 (c), TIGIT (d) and CD2 (e) is shown on CD32+ and CD32-CD4 T cells from HIV+ individuals (n=20) at 1 year following the initiation of antiretroviral therapy and controls (n=10). In panels b-d the percentage of cells expressing each marker is shown; for e the median fluorescence intensity (MFI) is shown. (f) Expression of H LA-DR on CD32+ and CD32-CD4 T cells from peripheral blood is shown from HIV+ individuals (n=8) at 1 year following the initiation of antiretroviral therapy. The expression of HIV co-receptors CCR5 (g) and CXCR4 (h) is shown between CD32+ and CD32-CD4 T cells from healthy controls. For CCR5 the percentage of cells expressing the marker is shown; for CXCR4 this is the MFI. Throughout, CD32+ cells (closed symbols) and CD32-cells (open symbols) were compared using the Wilcoxon matched-pairs signed rank test. Abbreviations used: ICR, immune checkpoint receptor; SSC, side-scattered light. ∗∗∗∗ indicates p<0.0001, ∗∗ indicates p 0.001-0.01, ∗ indicates p 0.01-0.05.

CD2 is ubiquitously expressed on CD4 T cells, but higher density of CD2 expression has been associated with enriched HIV DNA^10^. CD2 density (measured by median fluorescence intensity) was elevated on CD32+ compared to CD32-CD4 T cells (Figure 3e), consistent with these results.

### CD32+ CD4 T cells are activated and express high levels of HIV co-receptors

Amongst treated HIV-infected individuals (n=8, Supplementary Table 1) CD32+ CD4 T cells expressed markedly elevated levels of the activation marker HLA-DR (Figure 3f). In healthy controls, the expression of CCR5 and CXCR4, the co-receptors used by HIV for cellular entry, was increased on CD32+ CD4 T cells (Figure 3g,h).

### CD32+ CD4 T cells in gut and tonsil

CD32 expression on CD4 T cells in tonsil (HIV+ n=1), and terminal ileal and rectal biopsies (HIV+ n=1, control n=3) was at similar levels as in the periphery and did not appear to differ between HIV-infected and uninfected participants (Figure 4a). Sorted CD32+ CD4 T cells from tonsillar tissue were >4-fold enriched for HIV DNA relative to CD32-cells, and total CD4 T cells (0.025 vs 0.0061 and 0.0051 copies/cell respectively). Of note, this individual had commenced ART but was not virologically suppressed at the time of tonsillectomy (plasma VL 267 copies/ml). As in the periphery, CD32+ CD4 T cells from tissue were consistently more activated (as measured by HLA-DR expression) than CD32-cells (Figure 4b). Gut and tonsil CD32+ CD4 T cells showed a similar elevated pattern of ICR expression as in the periphery (Figure 4c), although there was considerable variation between biopsy sites.

**Figure 4.**
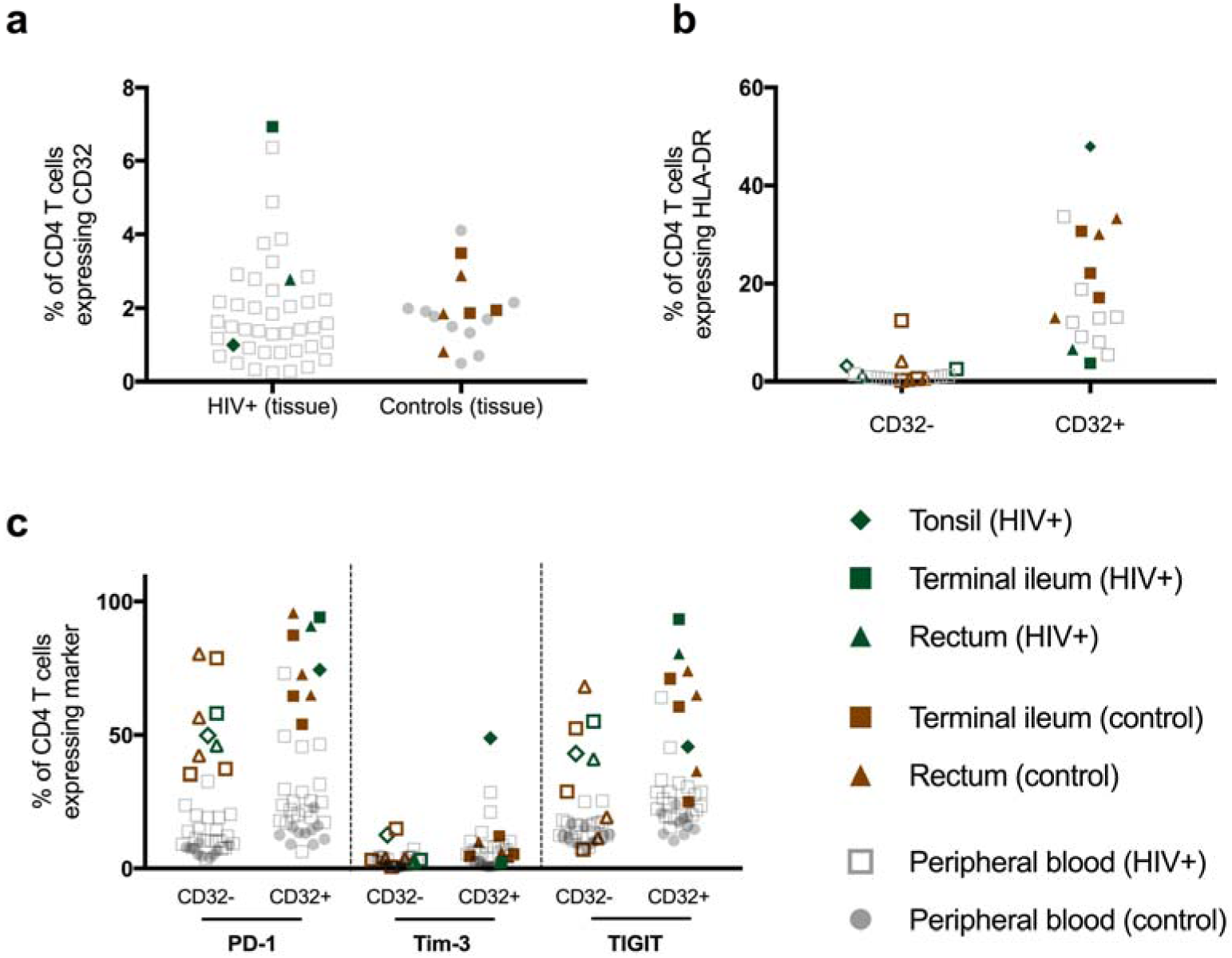
CD32 and immune checkpoint receptor expression in tissue. (a) The percentage of CD4 T cells expressing CD32 in tissue (HIV+ tonsil n=1, HIV+ gut n=1, HIV-gut n=3). (b) The expression of HLA-DR in tissue samples. (c) The expression of immune checkpoint receptors PD-1, Tim-3 and TIGIT on CD32+ (shaded shapes) and CD32-(open shapes) CD4 T cell populations in these tissues for HIV+ and HIV-individuals. Throughout, values obtained in tissue are superimposed on those from the periphery (HIV+ shown as open, grey squares and controls as shaded, grey circles) for visual comparison.

## Discussion

The report of CD32 as a marker of an enriched T cell HIV reservoir is potentially a key milestone in the search for a cure for HIV infection. To explore this finding further, we analysed individuals treated with ART early during PHI. We confirm that CD32+ CD4 T cells in peripheral blood are enriched for HIV DNA and provide the first data from tissue to support this association. We show that CD32+ CD4 T cells have a more differentiated memory phenotype, and express high levels of PD-1, Tim-3 and TIGIT, as well as the activation marker HLA-DR. Importantly, we show that the frequency and phenotype of CD32+ CD4 T cells is similar between uninfected and infected individuals - both prior to and following the initiation of ART. Our characterisation of CD32+ cells in tonsil and gut provides further insight into the relevance of these cells to the overall HIV reservoir *in vivo*.

Although the original association of CD32 expression with the HIV reservoir was unexpected, our findings are consistent with current understanding of the distribution of the HIV reservoir. We show that CD32+ CD4 T cells have a more differentiated memory phenotype, consistent with the fact that memory T cells comprise a large part of the HIV reservoir. It is interesting that central memory cells - the memory subset with the highest level of proviral DNA^9, 14^ - were less frequent amongst CD32 expressing cells. The expression of PD-1 and TIGIT has been reported to mark an enriched reservoir, although the fold enrichment in HIV DNA observed for these markers is substantially lower than that observed by us and others for CD32^8, 9, 12^. The elevated expression of ICRs on CD32+ CD4 T cells may partially explain the enrichment observed in previous studies with PD-1 and TIGIT. High level of CD2 expression was also previously identified as a maker of an enriched reservoir using an *in-vitro* model that also identified CD32 as a potential reservoir marker^10^. We observed an increased density of CD2 expression on CD32+ CD4 T cells, consistent with these results.

In our analysis of 19 individuals the percentage of CD32+ CD4 T cells did not predict time to pVL rebound following treatment interruption, although it is interesting that in Figure 1a the three individuals with post-treatment virological remission all had CD32 expression below the median. Delayed time to pVL rebound has been associated with lower HIV reservoir size^20, 22, 23^ and is an outcome measure in HIV curative intervention trials. In a large subanalysis of SPARTAC, HIV DNA levels, baseline pVL and CD4 count were predictive of time to viral rebound^20^ but this was not seen in this smaller subset so we cannot rule out that our negative findings here are a consequence of being underpowered.

In the paper by Descours *et al*., the addition of an integrase inhibitor into the *in-vitro* model reduced CD32 expression, suggesting that HIV integration resulted in CD32 upregulation^12^. Our finding of similar frequency and phenotype of CD32+ cells from HIV+ individuals and uninfected controls is therefore interesting. Although unstimulated CD4 T cells are not generally expected to express CD32^11^, we are not the first to observe low-level expression on CD4 T cells from healthy controls^24^. The similar level of CD32 expression between the HIV-infected (treated and untreated) and uninfected groups studied here and the lack of correlation between CD32+ CD4 T cell frequency and reservoir size suggests that infection or integration events may not be the primary driver of inter-individual variation in CD32 expression level, which may also be impacted by genetic and other host factors. Whilst several polymorphisms in the FcγRII genes have been identified to have functional implications in HIV immunity^25, 26^, whether these polymorphisms influence expression levels on CD4 T cells is unknown.

Several isoforms of CD32 exist, of which CD32a (FcγRIIa) and CD32b (FcγRIIb) are the most widely expressed. These two isoforms have an almost identical extracellular domain, but different intracellular signalling domains such that signalling in antigen presenting cells is activating through CD32b, but inhibitory through CD32a^11, 27^ In the *in-vitro* model used to identify CD32 as a marker of the reservoir, Descours *et al* note that FCGR2A - the gene encoding the CD32a isoform - was specifically upregulated, but not that of other isoforms^12^. The anti-CD32 antibody clone FUN-2 does not distinguish between different CD32 isoforms, and as this was used in the Descours publication and our work, HIV enrichment cannot yet be associated with a particular CD32 isoform.

The HIV reservoir is considered to be comprised of resting CD4 T cells - the elevated expression of the activation marker HLA-DR on CD32+ cells observed here is therefore potentially surprising. Activated CD4 T cells are preferentially infected with HIV but rapidly die as a result of this process, and the HIV reservoir is thought to form from cells that are infected whilst activated and then transition to a resting state^15, 28, 29^. Taken together, the high levels of HLA-DR, ICR expression and advanced differentiation suggest that CD32+ CD4 T cells have a history of activation - consistent with the current understanding of how the reservoir develops and is maintained.

The expression of CD32 on CD4 T cells remains controversial and a role for low-affinity Fc receptors on CD4 T cells has not been well established^30, 31, 32^. Although CD4 T cells are generally not considered to express CD32, expression may be induced by activation *in vitro*^33, 34^. Expression of CD32b on memory CD8 T cells in murine infection models^35^ is associated with reduced cytotoxicity and expansion, reversible on CD32b blockade^35^. This has interesting parallels with other co-inhibitory pathways, raising the possibility that CD32 isoforms may have a similar regulatory role on activated, antigen experienced CD4 T cells. This hypothesis may be linked to the enrichment of the HIV reservoir in CD32+ and ICR expressing CD4 T cells. Further work is needed to elucidate the specific mechanisms for enrichment of the HIV reservoir in these cells, but preferential infection or persistence of these cells is an alternative explanation that would be consistent with our findings.

In summary, we confirm that CD32+ CD4 T cells are enriched for HIV DNA and provide the first data showing similar enrichment in tissue. We show that CD32+ CD4 T cells have a more differentiated memory phenotype, and express high levels of PD-1, Tim-3 and TIGIT, as well as the activation marker HLA-DR. We show that the frequency and phenotype of CD32+ CD4 T cells is similar between uninfected and HIV-infected individuals - both prior to and following the initiation of ART. Even though possibly a marker for enrichment rather than a definitive biomarker of the reservoir, the identification of the role of CD32 raises the possibility of new diagnostic and therapeutic strategies within the HIV cure field.

## Methods

### Participants

Participants with PHI were recruited as part of the HEATHER (HIV Reservoir targeting with Early Antiretroviral Therapy) cohort. PHI was identified through one of the following criteria: (a) HIV-1 positive antibody test within 6 months of a HIV-1 negative antibody test, (b) HIV-1 antibody negative with positive PCR (or positive p24 Ag or viral load detectable), (c) RITA (recent incident assay test algorithm) assay result consistent with recent infection, (d) equivocal HIV-1 antibody test supported by a repeat test within 2 weeks showing a rising optical density and (e) having clinical manifestations of symptomatic HIV seroconversion illness supported by antigen positivity. For inclusion in the cohort, participants with identified PHI commenced ART within 3 months of diagnosis, and did not have co-infection with Hepatitis B or C. Date of seroconversion was estimated as the midpoint of the dates of the most recent negative or equivocal test and positive test (criteria a and d above), the date of the test (b and e) or 120 days prior to the test date (c, the recency period of this assay). For our study, cryopreserved PBMCs were used from the closest pre-therapy sample to seroconversion (baseline) and from a sample 9-15 months after commencement of ART (1 year). Terminal ileum and rectal samples were obtained from a subset of participants in HEATHER. Tonsil tissue was obtained through the Imperial College Communicable Disease Group Biobank from one HIV-infected individual with PHI undergoing routine tonsillectomy, 2 months after acquiring HIV. Terminal ileum and rectal samples from uninfected controls were obtained from individuals undergoing routine endoscopy from the Translational Gastroenterology Unit Oxford Tissue Biobank as part of the Oxford BRC-funded Oxford GI Illness Biobank Study.

Time to rebound analyses were conducted with a subset of participants from the SPARTAC (Short Pulse Antiretroviral Therapy at HIV Seroconversion) trial (EudraCT Number: 2004-000446-20). This was a multi-centre, randomised controlled trial of short course ART during PHI, the full design of which is described elsewhere^19^. The criteria used to define PHI in this trial are similar to above. In brief, this trial enrolled adults with PHI from 25 sites in Australia, Brazil, Ireland, Italy, South Africa, Spain, Uganda and the UK. Participants with PHI were randomised to receive either no immediate ART (standard of care), or 12 or 48 weeks of ART, after which they underwent a TI. The primary trial endpoint was a composite of CD4 T cell count less than 350 cells/μL or the initiation of long term ART for any reason. Cryopreserved PBMCs were used from participants who received 48 weeks of ART and were virologically suppressed to <400 copies/mL at this time. Participants were included based on sample availability at the time of TI, and date of seroconversion was estimated as calculated previously^7^.

### Ethics statement

Recruitment for and studies within the HEATHER cohort were approved by the West Midlands—South Birmingham Research Ethics Committee (reference 14/WM/1104).

The SPARTAC trial was approved by the following authorities: the Medicines and Healthcare products Regulatory Agency (UK), the Ministry of Health (Brazil), the Irish Medicines Board (Ireland), the Medicines Control Council (South Africa) and the Uganda National Council for Science and Technology (Uganda). It was also approved by the following ethics committees in the participating countries: the Central London Research Ethics Committee (UK), Hospital Universitário Clementino Fraga Filho Ethics in Research Committee (Brazil), the Clinical Research and Ethics Committee of Hospital Clinic in the province of Barcelona (Spain), the Adelaide and Meath Hospital Research Ethics Committee (Ireland), the University of Witwatersrand Human Research Ethics Committee, the University of Kwazulu-Natal Research Ethics Committee and the University of Cape Town Research Ethics Committee (South Africa), Uganda Virus Research Institute Science and ethics committee (Uganda), the Prince Charles Hospital Human Research Ethics Committee and St Vincent’s Hospital Human Research Ethics Committee (Australia) and the National Institute for Infectious Diseases Lazzaro Spallanzani, Institute Hospital and the Medical Research Ethics Committee, and the ethical committee of the Central Foundation of San Raffaele, MonteTabor (Italy).

All participants have given informed consent for their participation in these studies.

### Processing of tissue samples

Tonsillar tissue was dissected and mechanically digested, prior to cryopreservation of the cellular suspension.

Rectal and terminal ileum biopsies (up to 12 from each site) were collected at endoscopy and immediately place in RPMI-1640 media with 5% heat-inactivated fetal bovine serum (FBS), 0.04 mg/mL gentamicin, 100 IU/mL penicillin, 0.1 mg/mL streptomycin and 2mM L-glutamine. Biopsies were processed within 3 hours of sampling. Briefly, samples were washed in 1mM dithiothreitol (DTT) solution and then with PGA solution (Hanks’ Balance Salt Solution with 0.04 mg/mL gentamicin, 100 IU/ml penicillin and 0.1 mg/mL streptomycin). Biopsy samples subsequently underwent collagenase and mechanical digestion using Collagenase D (1 mg/mL) for 30 minutes and a gentleMACS dissociator (Miltenyi Biotec), respectively. The resulting cell suspension was then strained using a 70 μM filter, washed with PGA and used for staining.

### Flow cytometry and cell sorting

Cryopreserved PBMCs or tonsillar tissue were thawed in RPMI-1640 medium supplemented with 10% FBS, L-glutamine, penicillin and streptomycin as above (R10) containing 2.7 Kunitz units/mL of DNAse (Qiagen). For analysis of memory phenotype and ICR expression, cells were stained in BD Horizon Brilliant Stain Buffer (BD) containing all antibodies and Live/Dead Near IR at 1 in 300 dilution (Life Technologies) at 4°C for 30 minutes. PBMCs were stained with the following antibodies: CD32 PE-Cy7 (FUN-2), CD3 Brilliant Violet (BV) 570 (UCHT1), CCR7 Pacific Blue (G043H7), CD27 AlexaFluor700 (M-5271), CD2 APC (RPA-2.10)[BioLegend], CD4 BV 605 (RPA-T4), CD8 BV 650 (RPA-T8)[BD], PD-1 PE eFluor 610 (eBioJ105), CD45RA FITC (HI100), TIGIT PerCP-eFluor710 (MBSA43)[eBioscience] and Tim-3 PE (344823)[R&D]. Isotype controls for CD32 were prepared using an irrelevant IgG2bκ antibody (MPC-11)[BioLegend]. For time to rebound analyses, cells were stained as above in PBS with 5% FBS and 1mM EDTA containing Live/Dead Near IR, anti-CD32, anti-PD-1, anti-TIGIT as well as the following antibodies: CD3 FITC (UCHT1)[BioLegend] and CD4 eFluor450 (OKT4)[eBioscience]. Samples for HIV co-receptor stains were stained with LiveDead Near IR, anti-CD32 PE-Cy7, anti-CD3 FITC and anti-CD4 eFluor450 (as above). Following this, cells were incubated at 37°C for 20 minutes with anti-CCR5 PE (C57BL/6)[BD] or anti-CXCR4 APC (12G5)[BioLegend].

Mucosal cells were stained with LiveDead Near IR, anti-CD3 BV570, anti-CD4 BV605, anti-CD8 BV650 and anti-TIGIT PE-eFluor710 as above. The following antibodies were also included: PD-1 PE-Cy7 (EH12.1), Tim-3 PE-CF594 (7D3), HLA-DR AlexaFluor700 (MAB)[BD] and CD32 PE (FUN-2)[BioLegend]. EPCAM APC-Vio770 (HEA-125) [Miltenyi Biotec] was also used to allow for the exclusion of epithelial cells. Staining was performed for 20 minutes at room temperature, followed by fixation and permeabilisation using the Human FoxP3 Buffer Set (BD Pharmingen) as per manufacturers protocol.

All samples were acquired on a LSR II (BD). The same machine was used for all experiments with daily calibration with Rainbow Calibration Particles (BioLegend) to maximise comparability between days. Data were analysed using FlowJo Version 10.8.0r1 (Treestar).

For sorting experiments, CD4 T cells were enriched from cells thawed as above by negative magnetic selection using EasySep Human CD4+ T cell Enrichment kit (StemCell Technologies). CD4 T cells were stained in R10 at 4°C using Live/Dead Near IR, anti-CD32 PE-Cy7, anti-CD3 FITC, anti-CD4 eFluor450. Sorting of CD32+ and CD32-CD4 T cells was performed using a Mo-Flo XDP.

### Measurement of HIV DNA

For measurement of HIV DNA in bulk CD4 T cells, CD4 T cells were enriched from cryopreserved PBMCs as above or using Dynabeads Untouched Human CD4 T Cell Enrichment kit (Invitrogen). DNA was extracted from enriched CD4 T cells (QIAamp Blood Mini Kit; Qiagen) and sorted CD4 T cell subsets (QIAmp DNA Micro Kit, Qiagen or TRIzol extraction) and used as input for qPCR assays. Copies of HIV-1 DNA were quantified and normalised to number of input cells (as determined by albumin qPCR), by a previously described assay^20, 36^. Where possible, PCRs were performed in triplicate although due to the rare cell populations this was not possible for the sorted CD32+ cells.

### Statistical analysis

Continuous variables were compared between groups using non-parametric tests throughout. Comparisons between CD32+ and CD32-populations were performed using the Wilcoxon matched-pairs signed rank test. Where three groups were compared, a Kruskal-Wallis test (unpaired data) or Friedman test (paired data) was used; pairwise comparisons were performed on predetermined combinations of groups only if the overall test p-value was <0.05. Correlative analyses were performed using Spearman’s rank correlation. Time to viral rebound was assessed as time from treatment interruption to the first of two consecutive VL measurements >400 copies/mL (the limit of detection of the assay used at some trial sites). Individuals who did not rebound were censored at the date of the last VL measurement. Time to viral rebound was visualised with Kaplan-Meier curves stratified at the median and associations examined using Cox proportional hazard models. For all tests, p values <0.05 were considered statistically significant. Analyses were performed using GraphPad Prism (GraphPad Software, La Jolla, California, USA) version 6.0f or R version 3.2.2.

## Acknowledgments

We thank the participants of SPARTAC and HEATHER. The HEATHER study is conducted as part of the CHERUB (Collaborative HIV Eradication of Reservoirs: UK BRC) collaboration. (CHERUB Steering Committee: Andrew Lever (University of Cambridge), Mark Wills (University of Cambridge), Jonathan Weber (Imperial College, London), Sarah Fidler (Imperial College, London), John Frater (University of Oxford), Lucy Dorrell (University of Oxford), Mike Malim (King’s College, London), Julie Fox (King’s College London), Ravi Gupta (University College London), Clare Jolly (University College London).

The SPARTAC Trial Investigators: Trial Steering Committee: Independent Members-A Breckenridge (Chair), P Clayden, C Conlon, F Conradie, J Kaldor∗, F Maggiolo, F Ssali, Country Principal Investigators-DA Cooper, P Kaleebu, G Ramjee, M Schechter, G Tambussi, JM Miro, J Weber. Trial Physician: S Fidler. Trial Statistician: A Babiker. Data and Safety Monitoring Committee (DSMC): T Peto (Chair), A McLaren (in memoriam), V Beral, G Chene, J Hakim. Co-ordinating Trial Centre: Medical Research Council Clinical Trials Unit, London (A Babiker, K Porter, M Thomason, F Ewings, M Gabriel, D Johnson, K Thompson, A Cursley∗, K Donegan∗, E Fossey∗, P Kelleher∗, K Lee∗, B Murphy∗, D Nock∗). Central Immunology Laboratories and Repositories: The Peter Medawar Building for Pathogen Research, University of Oxford, UK (R Phillips, J Frater, L Ohm Laursen∗, N Robinson, P Goulder, H Brown). Central Virology Laboratories and Repositories: Jefferiss Trust Laboratories, Imperial College, London, UK (M McClure, D Bonsall∗, O Erlwein∗, A Helander∗, S Kaye, M Robinson, L Cook∗, G Adcock∗, P Ahmed∗). Clinical Endpoint Review Committee: N Paton, S Fidler. Investigators and Staff at Participating Sites: Australia: St Vincents Hospital, Sydney (A Kelleher), Northside Clinic, Melbourne (R Moore), East Sydney Doctors, Sydney (R McFarlane), Prahran Market Clinic, Melbourne (N Roth), Taylor Square Private Clinic, Sydney (R Finlayson), The Centre Clinic, Melbourne (B Kiem Tee), Sexual Health Centre, Melbourne (T Read), AIDS Medical Unit, Brisbane (M Kelly), Burwood Rd Practice, Sydney (N Doong), Holdsworth House Medical Practice, Sydney (M Bloch), Aids Research Initiative, Sydney (C Workman). Coordinating Centre in Australia: Kirby Institute University of New South Wales, Sydney (P Grey, DA Cooper, A Kelleher, M Law). Brazil: Projeto Praca Onze, Hospital Escola Sao Francisco de Assis, Universidade federal do Rio de Janeiro, Rio de Janeiro (M Schechter, P Gama, M Mercon∗, M Barbosa de Souza, C Beppu Yoshida, JR Grangeiro da Silva, A Sampaio Amaral, D Fernandes de Aguiar, M de Fatima Melo, R Quaresma Garrido). Italy: Ospedale San Raffaele, Milan (G Tambussi, S Nozza, M Pogliaghi, S Chiappetta, L Della Torre, E Gasparotto), Ospedale Lazzaro Spallanzani, Roma (G DOffizi, C Vlassi, A Corpolongo). South Africa: Cape Town: Desmond Tutu HIV-1 Centre, Institute of Infectious Diseases, Cape Town (R Wood, J Pitt, C Orrell, F Cilliers, R Croxford, K Middelkoop, LG Bekker, C Heiberg, J Aploon, N Killa, E Fielder, T Buhler). Johannesburg: The Wits Reproductive Health and HIV-1 Institute, University of Witswatersrand, Hillbrow Health Precinct, Johannesburg (H Rees, F Venter, T Palanee), Contract Laboratory Services, Johannesburg Hospital, Johannesburg (W Stevens, C Ingram, M Majam, M Papathanasopoulos). Kwazulu-Natal: HIV-1 Prevention Unit, Medical Research Council, Durban (G Ramjee, S Gappoo, J Moodley, A Premrajh, L Zako). Uganda: Medical Research Council/Uganda Virus Research Institute, Entebbe (H Grosskurth, A Kamali, P Kaleebu, U Bahemuka, J Mugisha∗, HF Njaj∗). Spain: Hospital Clinic-IDIBAPS, University of Barcelona, Barcelona (JM Miro, M Lopez-Dieguez∗, C Manzardo, J. Ambrosioni, D. Nicolas, JA Arnaiz, T Pumarola, M Plana, M Tuset, MC Ligero, MT Garca, T Gallart, JM Gatell). UK and Ireland: Royal Sussex County Hospital, Brighton (M Fisher, K Hobbs, N Perry, D Pao, D Maitland, L Heald), St James’s Hospital, Dublin (F Mulcahy, G Courtney, S O’Dea, D Reidy), Regional Infectious Diseases Unit, Western General Hospital and Genitourinary Dept, Royal Infirmary of Edinburgh, Edinburgh (C Leen, G Scott, L Ellis, S Morris, P Simmonds), Chelsea and Westminster Hospital, London (B Gazzard, D Hawkins, C Higgs), Homerton Hospital, London (J Anderson, S Mguni), Mortimer Market Centre, London (I Williams, N De Esteban, P Pellegrino, A Arenas-Pinto, D Cornforth∗, J Turner∗), North Middlesex Hospital (J Ainsworth, A Waters), Royal Free Hospital, London (M Johnson, S Kinloch, A Carroll, P Byrne, Z Cuthbertson), Barts & the London NHS Trust, London (C Orkin, J Hand, C De Souza), St Marys Hospital, London (J Weber, S Fidler, E Hamlyn, E Thomson∗, J Fox∗, K Legg, S Mullaney∗, A Winston, S Wilson, P Ambrose), Birmingham Heartlands Hospital, Birmingham (S Taylor, G Gilleran). Imperial College Trial Secretariat: S Keeling, A Becker. Imperial College DSMC Secretariat: C Boocock. (∗ Left the study team before the trial ended.)

We thank Satish Keshav, Carolina Arancibia (Translational Gastroenterology Unit, Oxford University Hospitals NHS Trust, as part of the Oxford BRC-funded Oxford GI Illness Biobank Study) and the participants who donated terminal ileum and rectal biopsies.

## Author Contributions

The experiments were conceived and designed by GM, MP, JT, CW, PKl, SF and JFr. Experiments were performed by GM, MP, JT, CP, MG, EH, HB, NR, NO and data analysed by GM, MP, JT, KP and JFr. Design and recruitment of the trials was performed by NN, JFo, SF, GR, PKa and JFr, with trial management performed by JM. The paper was written by GM and JFr with input from all authors.

